# Ancient Mongolian aurochs genomes reveal sustained introgression and management in East Asia

**DOI:** 10.1101/2023.08.10.552443

**Authors:** Katherine Brunson, Kelsey E. Witt, Susan Monge, Sloan Williams, David Peede, Davaakhuu Odsuren, Dashzeveg Bukhchuluun, Asa Cameron, Paul Szpak, Chunag Amartuvshin, William Honeychurch, Joshua Wright, Sarah Pleuger, Myagmar Erdene, Dashtseveg Tumen, Leland Rogers, Dorjpurev Khatanbaatar, Byambatseren Batdalai, Ganbaatar Galdan, Lisa Janz

**Author notes:** These authors contributed equally to this work (Katherine Brunson, Kelsey E. Witt).

## Abstract

Societies in East Asia have utilized domesticated cattle for over 5000 years, but the genetic history of cattle in East Asia remains understudied. Genome-wide analyses of 23 ancient Mongolian cattle reveal that East Asian aurochs and ancient East Asian taurine cattle are closely related, but neither are closely related to any modern East Asian breeds. We observe binary variation in aurochs diet throughout the early Neolithic, and genomic evidence shows millennia of sustained male-dominated introgression. We identify a unique connection between ancient Mongolian aurochs and the European Hereford breed. These results point to the likelihood of human management of aurochs in Northeast Asia prior to and during the initial adoption of taurine cattle pastoralism.

**One-Sentence Summary:** Ancient interbreeding of East Asian aurochs and cattle suggests management, but leaves no signature in modern eastern breeds.

## Main Text

Paleogenomic studies of cattle domestication largely focus on their origins in Western Eurasia, despite knowledge of multiple indigenous domestications of bovines such as yaks in the east. Cattle pastoralism has a long history in East Asia, but many questions remain about the origins of East Asian domestic cattle, including the possibility of indigenous domestication from local extinct wild aurochs (*Bos primigenius*). Current genetic evidence shows that modern cattle lineages largely derive from two aurochs domestication episodes: one in the Near East by 10,000 years ago that gave rise to taurine cattle (*Bos taurus*), and one in South Asia by 8000 years ago that gave rise to indicine/zebu cattle (*Bos indicus*) (*1-5*). Both genetic and archaeological evidence suggest the possibility of additional domestication events, most notably in North Africa (*6-9*), but the ability to conclusively recognize introgression over indigenous domestication remains largely unresolved. Regardless, wild introgression is acknowledged as critical to the development of modern lineages (*10-15*).

In East Asia today, taurine cattle breeds are more common in the north and indicine cattle breeds are more common in the south. There are many taurine-indicine hybrid breeds, and admixture between domestic cattle and other indigenous East Asian bovines such as yak, gayal/mithun, and banteng is common (*15-18*), with taurine-yak hybrids especially common in Central Mongolia. The north/south division between taurine and indicine cattle use appears to have deep roots related to the initial adoption of domestic cattle in East Asia. Taurine cattle were introduced to Northeast Asia at least 5000 years ago. Their bones are found at Afanasievo sites in Mongolia by about 5000 years ago (*19*), northeastern China by about 5300 years ago (*20*), and the Yellow River Valley by about 4500 years ago (*21*). Indicine cattle have played a minor role in Northeast Asia and are less well-understood, being present in southern China by the late Warring States period (ca. 500 BCE) (*22*). Here we explore the unique genetic history of aurochs and taurine cattle in Northeast Asia.

There is growing archaeological evidence that people in Northeast Asia hunted and possibly managed wild aurochs for thousands of years prior to the adoption of domestic taurine cattle. On the Mongolian steppe, sedentary societies during the mid-Holocene relied heavily on aurochs, with increasingly sedentary groups continuing to specialize in big-game hunting (*23, 24*). In northeastern China and the Yellow River Valley, several late Neolithic and early Bronze Age sites contain both aurochs and taurine cattle remains (*25, 26*). For example, at Houtaumuga in Jilin, archaeologists have uncovered pits full of aurochs bones. At this same site, geneticists have also identified the earliest East Asian cattle bone belonging to a taurine mtDNA haplogroup directly radiocarbon dated to 5500-5300 cal. BP (*20*). These finds indicate that there would have been opportunities for admixture between wild and domestic herds as soon as taurine cattle were first introduced to East Asia. However, only one East Asian aurochs mitogenome has been previously published (*27*) and there have not yet been any genome-wide ancient DNA studies of East Asian aurochs. Many questions remain, including how are East Asian aurochs populations related to East Asian domestic cattle? Were aurochs domesticated in East Asia? And did aurochs and domestic cattle interbreed when taurine cattle were first adopted in East Asia?

We sequenced low coverage genomes of 23 cattle excavated from seven archaeological sites in eastern Mongolia dated to between 30,000 years ago and 2,000 years ago (**Figure 1A; Table S1-S2**). The 30,000-year-old Paleolithic individual from Otson Tsokhio (*28*) represents the oldest paleogenomic data from an East Asian aurochs published to date. Early Neolithic samples from Tamsagbulag, Zaraa Uul, and Margal were directly radiocarbon dated to between 8,000-5,500 years ago and pre-date the introduction of taurine cattle. Bronze Age and later samples from Zaraa Uul (including the highest coverage sample B17), Delgerkhaan Uul, Shiriin Chuluu, and Ulaanzuukh date to after 4000 years ago. The genomes are low coverage (ranging from 0.01-0.94x), but they are still informative for investigating the deep history of cattle in Northeast Asia. We compared the ancient Mongolian samples with a reference panel of over 300 ancient and modern cattle genomes from around the world, including ancient taurine cattle from the 4000-year-old site of Shimao in the Ordos region of China (**Table S3**) (*5, 17, 29*). Shimao was one of the first sites in the Yellow River Valley region to use domesticated cattle in large numbers (*30, 31*). We also compared mitochondrial d-loop regions with previously published samples from China including the 6000-year-old individuals from Houtaomuga (*20*).

**Fig. 1.**
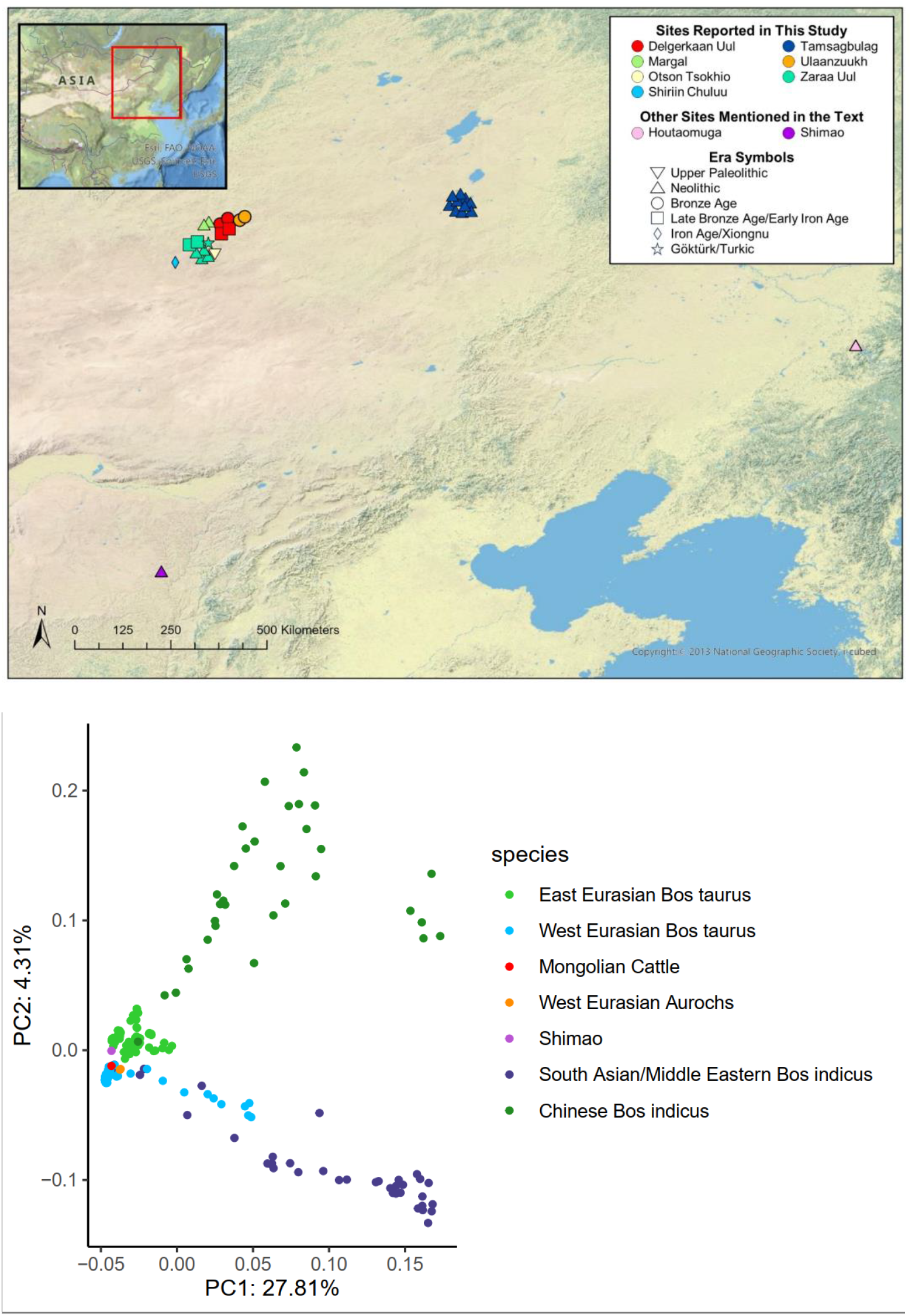
Geographical distributions and principal component analysis of ancient Mongolian cattle samples. (**A**) map showing the locations of samples according to time period. (**B**) PCA showing highest coverage ancient samples from Zaraa Uul (B17, early Bronze Age, labeled “Mongolian Cattle”) and Shimao (Shimao05, early Bronze Age) projected onto modern cattle individuals. High coverage ancient cattle individuals from Verdugo et al. (2019) are also included in the projection.

Mongolian aurochs show a close relationship to other sequenced aurochs and taurine cattle, and are distinct from indicine cattle (**Figure 1B**). The ancient Mongolian cattle form a single clade, suggesting that all sampled individuals represent a continuous population (**Figure 2B**). Neolithic Mongolian aurochs are part of the East Asian mitochondrial haplogroup C that has been previously identified exclusively in East Asian aurochs dating from between 10,000 to 4000 years ago (*20, 27, 32*) (**Figure 2A**). Interestingly, the Mongolian aurochs group together separately from African, European, and the Middle Eastern aurochs, but are still part of the clade that includes all sequenced aurochs and taurine cattle. Indicine cattle represent an outgroup. This suggests that aurochs across Northern Africa and Eurasia represent one large, panmictic population. Another possibility is that the Mongolian aurochs are part of the taurine cattle clade because they genetically contributed to early domesticated cattle in a significant way, although this hypothesis is difficult to test with low-coverage data.

**Fig. 2.**
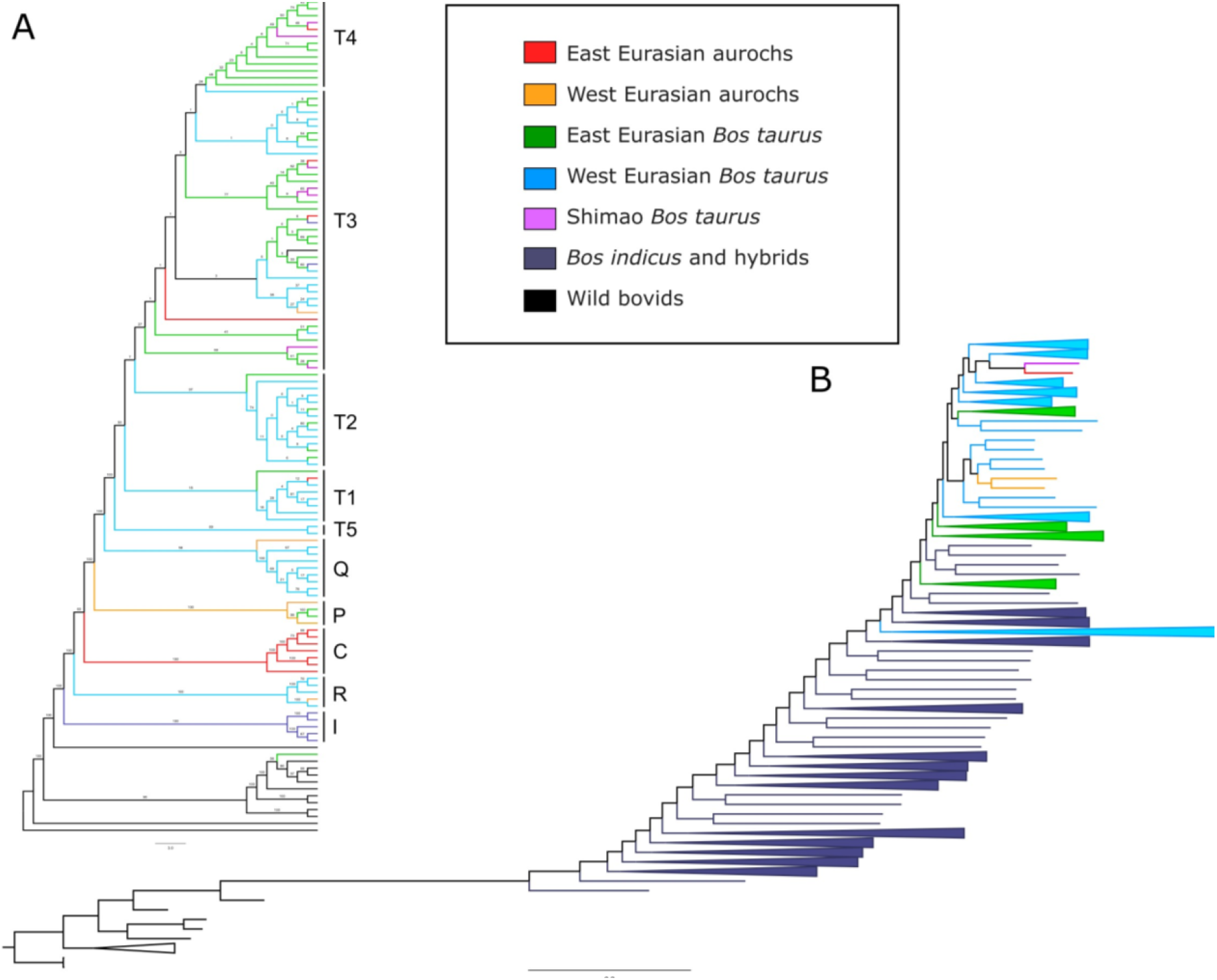
Phylogenetic relationships for cattle. (**A**) Maximum likelihood tree of whole mitochondrial genome sequences of modern and ancient cattle and bovids. (**B**) Maximum likelihood tree of genome-wide SNPs for all high-coverage bovids and ancient and modern cattle. Individuals are color-coded by geographic region and species. Hybrid cattle with any amount of indicine admixture are labeled as ‘*Bos indicus* and hybrids.’

The ancient Mongolian cattle are most closely related to 4000-year-old individuals from Shimao. Previous examination of modern Eurasian cattle breeds suggests three structured geographic groups: West Eurasian, Central Eurasian, and East Asian (*17*). The East Asian component of ancestry is found today primarily in Japanese cattle such as Mishima and was proposed to derive from gene flow between East Asian aurochs and domesticated taurine cattle. However, we do not observe any of this ancestry in ancient Mongolian aurochs, suggesting that another wild bovine population may have contributed to East Asian cattle (**Figure 3A**).

**Fig. 3.**
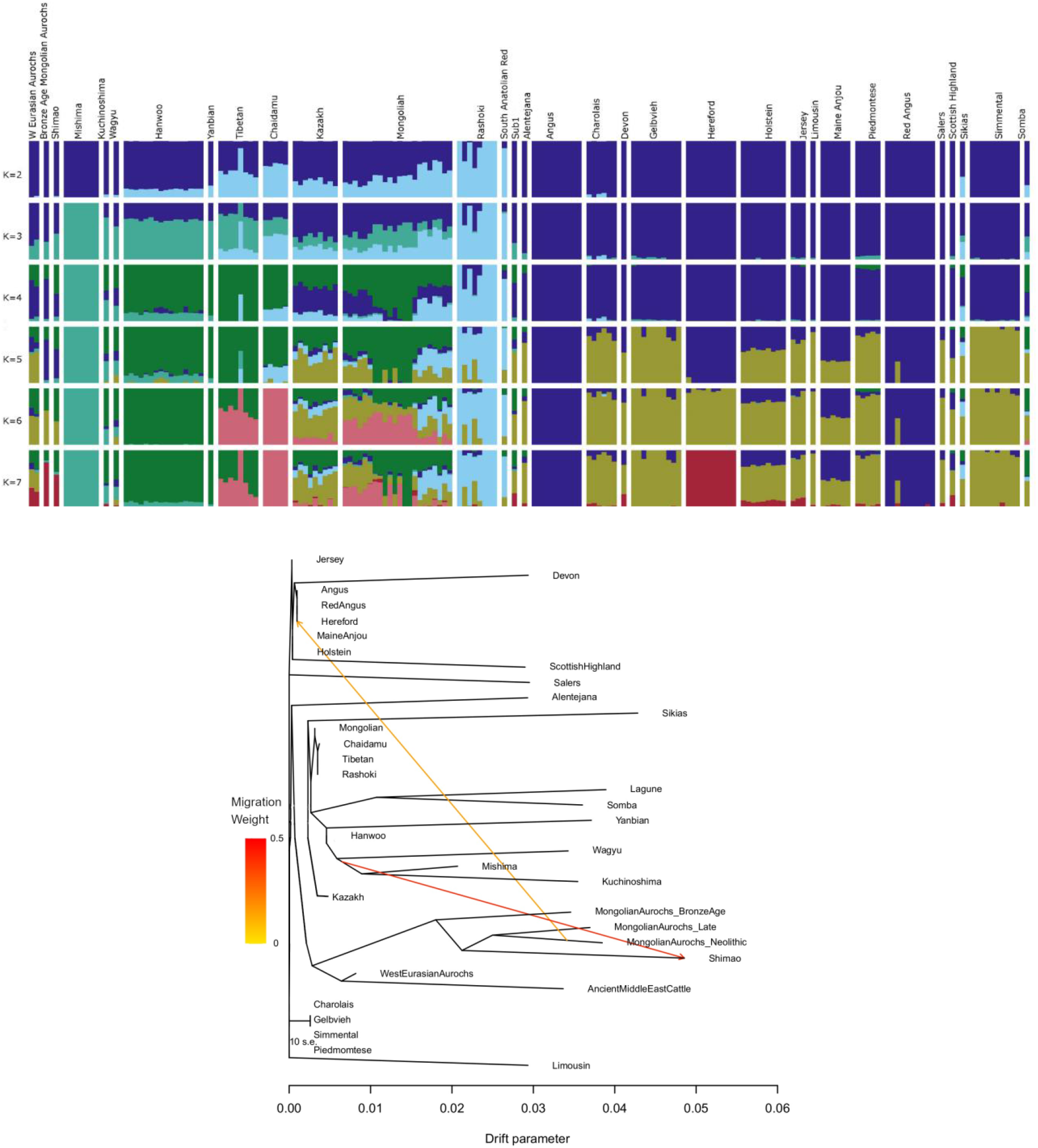
Admixture between Mongolian cattle and other populations. (**A**) Admixture plot showing population structure in taurine cattle and aurochs. At K=7 Mongolian aurochs, ancient Chinese cattle from Shimao, and Hereford have a shared ancestry component. (**B**) Treemix plot showing 2 migration edges among taurine cattle populations. Here, B01 is labeled as MongolianAurochs_Neolithic, B17 is MongolianAurochs_BronzeAge, and B28 is MongolianAurochs_Late.

Additionally, most modern cattle breeds seem to show very little to no ancestry from ancient Mongolian cattle. The one exception is Hereford, a British breed of beef cattle. Previous work has shown a connection between Hereford and cattle from Shimao (*17*), which are genetically similar to the ancient Mongolian cattle analyzed here. Our analyses show evidence for gene flow from Neolithic Mongolian aurochs into Hereford (**Figure 3B**). Herefords originated as traction and meat cattle from the long-isolated region of Herefordshire, England, where the cattle were heavily influenced by stock from the north of Wales but otherwise isolated until after 1700 (*33*). Perhaps the connection with ancient Mongolian cattle results from close genetic similarity between Mongolian aurochs and those of the British Isles, or perhaps both European and Mongolian aurochs contributed to early ancient taurine lineages, but their genetic legacy has since been swamped in other breeds. The connection might also be explained by more recent gene flow from a modern Asian cattle breed with Mongolian aurochs ancestry, but we find this hypothesis unlikely for two reasons. First, Mongolian aurochs ancestry is found almost exclusively in European breeds of cattle while modern cattle breeds from Mongolia and other parts of East Asia have little to no Mongolian aurochs ancestry (**Figure 3A**). Second, the gene flow from Mongolian aurochs seems to come specifically from Neolithic individuals, which pre-date the introduction of taurine cattle to East Asia (**Figure 3B**). Additional sequencing of ancient cattle from across Eurasia would be needed to clarify this connection.

Our analysis also reveals one additional admixture event from Japanese cattle breeds into Shimao (**Figure 3B**). The gene flow from East Asian cattle lineages into Shimao is consistent with previous work that shows a connection between ancient Shimao cattle and modern northeast Asian breeds (*17*). Cattle from Shimao form a clade with the ancient Mongolian cattle and also have taurine mitochondrial haplogroups, suggesting hybridization between East Asian aurochs and taurine cattle. All individuals predating the arrival of taurine cattle are from mitochondrial haplogroup C. While haplogroup P, which is found in European aurochs, has been identified in modern Japanese cattle (*34*), we did not observe haplogroup P in the Neolithic Mongolian aurochs. Most later individuals are mitochondrial haplogroup T3, which is most common in both East Asia and western Eurasia. One individual (B20) is haplogroup T1, which is common today only in southern Europe and Africa (*35*). One individual (B12) from the sixth century is Haplogroup T4, but still bears significant evidence of Mongolian aurochs ancestry. The T4 haplogroup is distinct to East Asian and believed to have diverged locally from earlier arrivals of T3 or admixture with East Asian aurochs (*10, 36*). We do not think that the introduction of taurine cattle represents population replacement because local Bronze Age cattle are still very similar to Neolithic aurochs and genetically distinct from ancient western Eurasian cattle (*5*). These preliminary findings suggest that early gene flow was primarily between male aurochs and female taurine cattle. The lack of modern cattle with aurochs DNA suggests that these aurochs lineages were replaced over time, likely due to interbreeding with other Eurasian cattle breeds. Aurochs persisted in China into the Zhou Dynasty (1046-256 BCE) and probably much later (*37*). More geographically and temporally extensive genomic analyses of ancient East Asian cattle would provide additional insights into how East Asian cattle populations changed through time.

Stable isotope analysis was carried out on 7 of the samples, of which 4 Neolithic individuals (B06, B13, B15, B23) had elevated δ13C and δ15N ratios locally consistent with foddering (*38*) (**Table S1**). Elevated δ13C and δ15N is observed to be characteristic of Late Neolithic aurochs at the Honghe site in Northeast China (*26*). C4 plants are not common in these predominantly C3 landscapes and our findings point to human intervention in diet. To test the possibility of aurochs management, we looked at measures of heterozygosity (**Figure S8**) in order to determine whether Mongolian individuals show changes in genetic diversity through time. A reduction in genetic diversity might be evidence for bottlenecks during management or domestication. Although heterozygosity may be slightly lower in the Bronze Age individuals compared to Neolithic individuals, we also see that genetic diversity remains high in some later samples such as the Xiongnu individual from Shiriin Chuluu. We do not observe a consistent trend in heterozygosity that shows direct evidence for aurochs management, nor is there an overlap between individuals that may have been foddered and individuals with taurine mitochondrial haplotypes. Individuals that may have been foddered are all early Neolithic (Oasis 2, 8500-5000 cal BP) aurochs and tend to group more closely in our phylogenetic tree regardless of age or geographic region (**Figure S9**). The results point to variation in aurochs diet where some pre-taurine individuals were being grazed on or foddered with C4 plants while others were uncontrolled grazers. Deeper coverage genomic and additional isotope analyses are needed to confirm whether elevated δ13C and δ15N is connected to distinct lineages and how that may or may not have related to later interbreeding. Our current results suggest that diets of the first taurine herds were similar to ones associated with open grazing practices followed by modern local herders. More broadly, our results show the persistence of aurochs ancestry through time, even in taurine cattle from the Xiongnu and Turkic periods, which suggests widespread intentional interbreeding. This goes beyond what we might expect from opportunistic mating with wild bulls during the early stages of introduction or periods of stress (*12*). The isotopic results support the possibility of indigenous aurochs management in Mongolia by ∼7800 cal BP, with genetic evidence for interbreeding between aurochs and taurine herds by ∼2900 cal BP.

Management of herd animals in East Asia is often viewed as an external process where fully domesticated cattle, caprines, and horses were introduced to eastern Siberia, Mongolia, and China around the same time that the first pastoralist groups arrived from western Eurasia (*19, 39-42*). East Asia has not traditionally been recognized as a center for cattle domestication. Our results instead suggest that humans were likely already managing aurochs before the arrival of taurine cattle to Northeast Asia around 5000 years ago, after which they began intentionally interbreeding the two populations. The earliest taurine cattle introduced to East Asia are known to have been used in dairying (*43*), which highlights the importance of female taurine lineages. Male based introgression in cattle has been previously posited for Eurasia and Africa (*9, 12, 17*), but the unreliability of Y-chromosome data and a lack of genome-wide studies for ancient samples has obfuscated the evidence. Previous work has underscored the importance of introgression in environmental adaptation during adoption of domesticates (*5, 12, 15*). Based on the dominance of domesticated taurine mtDNA, the importance of dairying in this region, and the sustained nature of interbreeding in our sample, we hypothesize that in Mongolia this process is more accurately characterized as the introgression of female dairy cattle into indigenous herds in order to facilitate milk production. Attempts to improve fecundity and offspring survivorship might also have played a role. A greater emphasis on controlling female lineages for milk production would also explain the Eurasian expansion of haplogroup T3 ∼7.5kyBP (*12*) after the emergence of dairying in the Near East (*44*).

Zooarchaeological and paleogenetic research increasingly shows that humans have managed animals in a variety of ways that do not always match the expectations of traditional models of animal domestication (*45-48*). Domestic animal populations often originate through multilocal processes, and admixture between wild and domestic populations was common for many taxa (*5, 49-51*). Our results show that these insights apply to cattle domestication in East Asia as well. Archaeological research focused on the presence and use of aurochs in and around habitation sites, as well as their individual life histories is ongoing and this work can add to our understanding of hunter-gatherer interactions with aurochs in Mongolia and also broader questions of aurochs management across East Asia.

The expansion of cattle pastoralism in East Asia intensified after about 4000 years ago. This coincided with broader changes in subsistence, ritual practices, and landscape use. Archaeological finds of dairy products (*52*) and identification of milk residues in pottery and human dental calculus suggests that milk was important to Bronze Age and later pastoralist peoples across the mountain, steppe, and desert regions of East Asia (*43, 53, 54*). Cattle bones were heavily utilized for oracle bone divination in ancient China by 4000 years ago (*55*), and cattle were also used in large numbers for subsistence, bone artifact production, and ritual sacrifice at early urban centers by 3000 years ago (*56*). Increasing reliance on cattle coincides with a decline in wild fauna such as deer, suggesting changes in human landscape use away from the mosaic habitats that facilitated interactions between humans and wild fauna (*57*). We do not know when during these transitions wild aurochs went extinct, but our results suggest that we should not ignore the possibility that people actively managed aurochs both before and long after the introduction of taurine cattle. The history of domesticated cattle in East Asia from initial domestication to the present remains enigmatic and further study of the archaeological and genetic history of ancient populations will be critical for understanding aurochs management and how they contributed to early cattle husbandry and modern breeds.

## Acknowledgments

We thank the Gobi-Steppe Neolithic Project, Mongolian National University of Education, Mongolian Academy of Sciences, Mongolian National University, University of Illinois, Chicago sequencing and research informatics cores, Rush University Medical Center Genomics and Microbiome Core Facility, Trent University, and the Wesleyan high powered computing cluster. Special thanks to Sohini Ramachandran and the Ramachandran lab, Emelia Huerta-Sanchez and the Huerta-Sanchez lab, and the Brown Center for Computational Molecular Biology for their support and insights during the research for this project. Part of this research was conducted using computational resources and services at the Center for Computation and Visualization, Brown University. We thank Guy Bennevat Haninovich for assistance compiling modern reference genomes, Nina Hirai for assistance producing the map in Figure 1, and Henk Meij for assistance with the Wesleyan HPCC, Canyang Ye for assistance researching bovine use in the Chinese-language literature, and Adiyasuren Molor for help with Mongolian-English translation.

## Author contributions

Conceptualization: LJ, KB, KW

Methodology: KB, KW, SW, SM, LJ

Investigation: KB, KW, SW, SM, LJ, DO, DB, DP, LR

Visualization: KB, KW, DP

Funding acquisition: LJ

Writing–original draft: KB, KW, LJ, SW, SM, DP

Writing–review & editing: all contributing authors

## Data and materials availability

All raw reads from the project are available on the NCBI SRA archive at [links will be added at time of publication]; code is available on Github [links will be added at time of publication]; MtDNA alignments are available at [links will be added at time of publication].

