## Supplemental Figures for "Ancient Mongolian aurochs genomes reveal sustained introgression and management in East Asia"

### Materials and Methods

Archaeological Contexts

Our samples originate from archaeological sites in eastern Mongolia. **Table S1** lists contextual information for the 23 samples in this study including excavation context, skeletal elements represented, and dates. It also includes information for an additional 4 samples that we were unable to extract DNA from. In this section, we provide background on the archaeological sites and contexts of the samples.

Tamsagbulag is located in the far eastern steppe, along a terrace of a former riverbed. Delgerkhaan Uul, Margal, Ulaanzuukh, Shiriin Chuluu, Zaraa Uul and Otson Tsokhio are located in the desert-steppe region, all within a 100 km radius. Margal is an in situ Neolithic habitation site in the broader Delgerkhaan Uul region and dates to the end of Oasis 2, post-dating Zaraa Uul and Tamsagbulag by ~2000 years and pre-dating by a few centuries the earliest evidence for *Bos taurus* in East Asia (*20*).  Margal was excavated under the Mongol-American DMS survey project (58). Samples from Delgerkhaan Uul, Ulaanzukh (type site for the eponymous Bronze Age burial culture – see (59)) and Shiriin Chuluu were all excavated from burial contexts. Prone burials are the earliest form of burials associated with local pastoralist Bronze Age cultures, beginning ~3500 cal BP and lasting until ~3100 cal BP (*60*). Slab burials replace the prone burial types at the end of the Bronze Age and overlap with the beginning of the Iron Age, a period which culminates in the formation of the Xiongnu/Hunnu Empire in the 3rd century BC. Many of these burials include domesticated herd animals assumed to derive from the descendants of introduced Western lineages. Animal offerings are sometimes intrusive, having been placed in later periods (see B19 in Table S1 “Context”). The Otson Tsokhio habitation site spans the Initial and Early Upper Palaeolithic (IUP/EUP) and is located within 1 km of Zaraa Uul (28).

The majority of our samples come from Zaraa Uul and Tamsagbulag. Samples from Zaraa Uul span ~8500-1500 BP and include habitation and burial contexts. The earliest contexts are represented by a recurrent Early Neolithic/Oasis 2 occupation straddling the downslope of a heavily eroded mountain range and the shore of an expansive freshwater marsh. Neolithic components were secondarily deposited just downslope from their original context, and intermixed with a few materials dating to the Late Bronze Age, Early Iron Age, and Turkic periods. All such samples were directly dated due to the possibility of intermixing.

The site of Tamsagbulag is known for its intensity of aurochs exploitation, which led the original excavators to describe it as an agropastoralist campsite, belonging to sedentary, millet-farming cattle pastoralists (*61*, *62*). There is no evidence of millet or other farming activities and radiocarbon dates now show the site significantly predates the arrival of domesticated cattle in East Asia (*63*). Intensity of occupation is substantially higher than in the desert-steppe and is comparable to contemporary sedentary sites in the Inner Mongolia Autonomous Region, PRC (*24*). Deposits of highly organic soils excavated in 2018, show accumulations of up to ~80 cm within little more than 100 years. Substantial site architecture, including postholes, trench features, clay-lined hearths, and deep ash pits underscore the high intensity of occupation. Numerous radiocarbon dates for Tamsagbulag indicate that the site was used 8.4-6.0 ka cal BP, and most intensively at 7.8-7.5 ka cal BP (*24*).

Radiocarbon Dating

Dates for individual samples, where available, are reported in **Table S1**. Fourteen samples were dated directly using AMS 14C dating. Remaining samples were dated based on stratigraphic association. Samples for AMS 14C dating were selected from faunal remains based on their context. Bone collagen for all samples was extracted and purified at the Trent Environmental Archaeology Laboratory (TEAL) using Centriprep 30 kDa centrifugal ultrafilters according to the UCI-KCCAMS procedure (*64*). The samples were prepared alongside a woolly mammoth bone with an infinite radiocarbon age (Hollis Mine Mammoth, Fmc=0.0031±0.0009) and two secondary standards that have been measured multiple times (Umingmak Wale, Fmc=0.4011±0.0009; Banks Island Musk Ox, Fmc=0.6489±0.0011). The >30 kDa collagen extracts were sent to the A. E. Lalonde AMS Laboratory where radiocarbon analyses were performed on a 3MV tandem accelerator mass spectrometer built by High Voltage Engineering (HVE). 12,13,14C+3 ions were measured at 2.5 MV terminal voltage with Ar stripping (*65*). The fraction modern carbon, F14C, was calculated according to Reimer et al. (2004) as the ratio of the sample 14C/12C ratio to the standard 14C/12C ratio (in our case Ox-II) measured in the same data block. Both 14C/12C ratios were background-corrected and the result was corrected for spectrometer and preparation fractionation using the AMS measured 13C/12C ratio and normalized to δ13C (PDB). Radiocarbon ages were calculated as -8033ln(F14C) and reported in 14C yr BP (BP=AD 1950) as described by Stuiver and Polach 1977 (*67*). Calibration was performed using OxCal v4.4.2 (*68*) and the IntCal20 calibration curve (*69*).

Isotope Extraction

Bone collagen was extracted from 7 of the samples included in this study (**Table S1**). Briefly, bone (250-750 mg) was mechanically cleaned of foreign matter with a dental drill equipped with a diamond-tipped cutting wheel. The samples were demineralized in 0.5 M HCl at room temperature for several days, leaving an insoluble collagen residue, which was rinsed to neutrality with Type I water, then treated with 0.1 M NaOH for ~20 min between one and five times to remove humic contaminants. The samples were rinsed to neutrality with Type I water and then heated in sealed glass tubes at 75oC for 16 h in 0.01 M HCl to hydrolyze the collagen, which was transferred into pre-weighed glass vials, frozen, and lyophilized. The carbon and nitrogen isotopic and elemental compositions of the collagen were determined using a Nu Horizon isotope ratio mass spectrometer coupled to a EuroVector 3300 elemental analyzer. The δ13C and δ15N values were calibrated relative to AIR and VPDB, respectively using USGS40 and USGS41a. In house reference materials were analyzed alongside the samples to monitor accuracy and precision: SRM-1 (caribou bone collagen), SRM-14 (polar bear bone collagen), SRM-20 (hydroxyproline). Thirty-five of the 123 unique tooth increment samples were analyzed in duplicate or triplicate. The overall analytical uncertainty was estimated to be ±0.19 for δ13C and ±0.30 for δ15N based on Szpak et al. 2017 (*70*).

### Ancient DNA Extraction and Sequencing

#### Sample preparation

A total of 29 bone or tooth samples were analyzed at the Williams Ancient DNA Laboratory with HEPA filtered positive airflow at the University of Illinois at Chicago (UIC), which is used exclusively for ancient DNA research and physically separated from the post-PCR laboratory. Rigorous contamination controls followed standard procedures, including multiple negative controls at all stages of analysis, the sterilization of surfaces and instruments by UV light, and the decontamination of the work area and equipment with 10% sodium hypochlorite solution. Before the DNA extraction, samples were decontaminated by submerging them in a 10% sodium hypochlorite solution for 4 minutes and subsequently rinsed in DNAse free water on a tube rocker for 15 minutes, repeating the rinse if necessary or until the water was clear.

#### DNA Extraction

The DNA extraction followed a modified silica-spin column method (*71*). The decontaminated samples were ground into a powder and approximately 120mg of bone powder was incubated for at least 18 hours at 37°C in lysis buffer (0.45M EDTA pH 8.0, 20mg/20mL Proteinase K). Afterwards, 1mL of sample supernatant was transferred to a Zymo-Spin V Extender Assembly along with 13mL of binding buffer (guanidine hydrochloride, sodium acetate, Tween-20, and silica suspension). The extracts were purified using QIAGEN PE buffer and eluted with QIAGEN Buffer EB. Each sample was extracted several times along with no more than 6 samples at a time and a negative control. DNA quantity was measured using a Qubit quantification platform. As a further check, extracted DNA samples were amplified and visualized on acrylamide gels in a laboratory located in another building on campus (*72*). Two sets of primers that target cattle mitochondrial D-loop control regions were used (L16022/H16178 and L16137/H16315 (*16*, *25*). Six samples did not show evidence of DNA and received no further analysis.

#### Next-Generation Sequencing

Genomic libraries were prepared for next-generation sequencing using the NEBNext® Ultra™ II DNA Library Prep kit for Illumina® with NEBNext® Multiplex Oligos (Dual Index primers) and purified using AMPure XP magnetic beads (Beckman Coulter). The NEBNext® Ultra™ II protocol was followed, with adaptor dilution at 1:15 and cleanup of adaptor-ligated DNA was carried out without size selection. The binding time with the beads was extended to 10 minutes. PCR amplification of the libraries included 14 cycles of denaturation at 98°C for 10 seconds and annealing/extension at 65°C for 75 seconds. Thirteen libraries were sequenced at the Genome Research Core at UIC, an additional 10 samples were sequenced at the Rush University Medical Center Genomics and Microbiome Core Facility. The library fragments were visualized on D1000 and D5000 ScreenTape Systems in both labs. An initial run was performed on an Illumina® MiniSeq to assess the number of reads that mapped to cattle DNA sequence. Quality assessment was undertaken by the UIC Research Bioinformatics Core Facility. Cattle read percentages ranged from 14-97%, with an average of 43%. Two samples (B1 and B17) were selected for deeper coverage sequencing because of their high quality. The highest cattle percentage (97% DNA reads) came from B17, the only petrous bone sample collected. An llumina® NovaSeq was used to generate 2X150 paired-end reads.

### Alignment to Reference Cattle Genome

All raw reads were analyzed using a standard pipeline optimized for ancient samples. Adapters were removed using AdapterRemoval v2 (*73*) and were aligned to a reference genome that combined the autosomes and X chromosome from BosTau6 (UCSC UMD3.1), the Y chromosome from BosTau7, and the mitochondrial reference sequence (accession V00654.1). The Burrows-Wheeler Alignment Tool version 0.7.15 with the 'mem' function (*74*) was used to align the reads to the reference. The resulting sam files were converted to bam files using the SAMtools version 1.9 view command, which removed unaligned reads and reads with a mapping quality less than 30 (samtools view -b -F 4 -q 30) (*75*). SAMtools was also used to sort the reads and remove duplicates (samtools sort and samtools rmdup, respectively). **Table S2** lists total genome coverage, number of nucleotides, and average coverage for each sample.

All comparative samples downloaded from the Sequence Read Archive (**Table S3**) as fastq files were aligned using the same pipeline, starting at the "bwa mem" step. The samples utilized from Verdugo et al. 2019 (the gaur, all other aurochs, and the ancient Near Eastern cattle) were used in the bam file format, as they were mapped to the same reference genome.

### Quality Control of NGS Data

We used DamageProfiler (*76*) to authenticate ancient sequences. Typical ancient DNA damage patterns have parabolic distributions with more C to T misincorporations at the 5 prime ends of reads and more G to A misincorporations at the 3 prime ends of reads. DamageProfiler deamination curves and length distribution results are shown in **Figures S1 and S2**.

### SNP Calling

The program ANGSD ver. 0.930 (77) was used to call variants. The method uses a genotype likelihood method, which accounts for some of the uncertainty surrounding ancient DNA SNP calls. Genotype likelihoods were called using the SAMtools model (-GL 1) for all sites with a minimum number of individuals represented (-minInd ###) where the number of individuals is equal to the number of modern samples plus half the number of ancient individuals, producing a Beagle file output (-doGlf 2). The major allele was polarized to the reference genome and allele frequencies for each allele were calculated (-doMajorMinor 1 -doMaf 2). A python script was used to call haploid genomes for each modern and ancient individual - transitions were filtered out and the base pair represented in the majority of reads were called for that individual (78). We used multiple subsets of the data depending on the analysis. "High Coverage Bovines" includes all Bos individuals (B. taurus, B. indicus, B. primigenius, B. frontalis, B. grunniens, B. gaurus, B. javanicus, B. mutus), wisent (Bison bonasus), and water buffalo (Bubalus bubalus), all modern cattle samples, and all ancient samples with at least 1x coverage, plus B17 from this publication. We also used a Taurus-Indicus dataset that consisted of the High Coverage Bovines that are part of B. indicus, B. taurus, and B. primigenius, and a Taurine dataset with only B. taurus and B. primigenius. For some analyses we included lower-coverage individuals in the dataset, and in these cases we down-sampled the high coverage data to only include the sites that were found in the low coverage Mongolian cattle. We also made a “Mongolian Cattle” dataset that consists of fifteen Mongolian cattle individuals and one European aurochs as outgroup; that dataset is limited to sites that are shared across all samples.

### Gene Tree

A whole-genome phylogeny was constructed using the SNP data for the “High Coverage Bovines” set. The vcf file was converted to a phylip alignment using a python script called “vcf2phylip.py” (https://github.com/edgardomortiz/vcf2phylip/blob/master/vcf2phylip.py) and a maximum likelihood tree was created using RAxML (79) using a rapid bootstrap analysis method (-f a) with 1000 replicates (-# 1000). We also made a gene tree of the Mongolian Cattle dataset downsampled to shared sites shown in **Figure S9**.

### Principal Component Analysis

We used smartpca from eigensoft v. 6.0 (80) to create a Principal Component Analysis. A python script was used to convert the vcf to the input files for smartPCA, and high-coverage ancient and modern individuals were used in the calculation of the eigenvectors, while the low-coverage ancient individuals (from this study and others) were projected onto the PCA (lsqproject: YES in the parameter file). ADMIXTURE version 1.3.0 (81) was used to construct admixture plots from k=2 to k=5. The PCA plot was conducted for the “High Coverage Bovines” set.

### Introgression Scans (D Statistics)

To test for the presence of Auroch introgression in Taurine cattle we utilized the D statistic which uses site patterns as a proxy for genealogical relationships to assess if there are more discordant topologies than expected under neutrality. Specifically we split all Taurine cattle by geographic groups, computed the D statistic for all trios for all pairwise combinations of Taurine geographic groups using the following configuration: P1 = Taurine individual, P2 = Taurine individual, P3 = Auroch, and P4 = Gaur, and lastly we assessed significance by using a standard block-jackniffing bootstrapping procedure as described in Durand et al. (2011) with a block size 50kb, which was determined based on known patterns of LD-decay in cattle following Verdugo et al. (2019). As was done in Verdugo et al. (2019), we filtered out any configurations that did not have at least 200 ABBA+BABA sites and considered a configuration statistically significant if the absolute value of the Z-score was greater than or equal to three. D statistics results are shown in **Table S7 and Figure S3**.

### Admixture Graphs

We used the “convertf” function in eigensoft version 6.0 (*80*) to convert the smartPCA input files to a format compatible with the program admixture. We used ADMIXTURE version 1.3.0 (*81*) to examine population structure within the Taurine dataset for K-values 2-7. Visualization was performed using the R script AdmixturePlotter (https://github.com/TCLamnidis/AdmixturePlotter). **Figure S4** shows a plot of three of the highest coverage ancient Mongolian samples.

### TreeMix

TreeMix version 1.13 (*83*) was used to create a TreeMix plot using the High Coverage Bovines to identify possible sources of gene flow (**Figure S5**). We simulated 0-5 migration edges, used *B. bison* as an outgroup population and performed 1000 replicates. We also performed TreeMix analyses for the downsampled datasets to see how the aurochs from different sites might affect gene flow.

### Mitochondrial DNA Analyses

We used SAMtools (*75*) to extract complete mitochondrial genomes from the sample bam files and then made an alignment using MUSCLE in MEGA11 (*84*). Samples B06, B07, B10, B11, B14, B15, B16, B19, B21, B23, B26, and B29 were not included in the analysis due to low coverage of the mitochondrial genome. Previously published mitochondrial genomes from modern and ancient cattle worldwide were added to the alignment (**Table S4)**.

We constructed a maximum likelihood tree for the complete mitochondrial genomes using RAxML v8.2.12 (*79*). The alignment was converted into phylip format and analyzed using rapid bootstrapping and searching for the best scoring ML tree, GTR + GAMMA model of rate heterogeneity and correcting for ascertainment bias, water buffalo NC049568 as outgroup, and we used 1000 bootstraps for replications. RAxML parameters were run as follows:

```
raxmlHPC-AVX2 -f a -n result -m GTRGAMMAX -p 12345 -N 1000 -k -x 12345 -o NC049568
-s alignmentinfile.phy
```

The resulting ML tree (**Figure S6**) reveals that all of the Neolithic samples (B01, B02, B03, B04, B05, and B13) are part of the C haplogroup of East Asian aurochs. The post-Neolithic samples (B12, B17, B20, B27, and B28) are part of the T1, T3, and T4 haplogroups of taurine cattle.

To further investigate variation among the ancient individuals, we identified variable nucleotide positions in part of the mtDNA d-loop region (16022-16315) as shown in **Table S5**. We also created a maximum likelihood phylogenetic tree for this d-loop region, excluding ancient Mongolian individuals with low coverage for the d-loop (**Table S6**). Individuals in the network include B01, B02, B03, B04, B05, B12, B13, B17, and B28. The network also includes most of the modern and ancient reference individuals from the complete mitogenome analysis plus additional previously published d-loop sequences from ancient individuals at Houtaomuga, which include a large number of individuals in the C haplogroup as well as the earliest T3 haplogroup identified in China (20). It also includes some modern Japanese Shorthorn breed individuals in the P haplogroup (34), and one additional ancient Mesolithic/Neolithic European aurochs individual in the P haplogroup (*85*). The ML tree was constructed in RaxML with the same parameters as listed above and with water buffalo NC006295 as an outgroup.

We used PopArt (*86*) to construct a median joining network for the d-loop alignment (**Figure S7**). The d-loop analysis is consistent with previous studies of East Asian bovines, showing that aurochs haplogroups are distinct from taurine cattle in the T haplogroups.

Heterozygosity

Heterozygosity was calculated for each individual in the High Coverage Bovines dataset to assess how the genetic diversity in the East Asian aurochs compared to other wild and domesticated bovine populations. Angsd and realSFS were used to construct site frequency spectra for each individual and calculate genome-wide heterozygosity. Results are shown in **Figure S8**.

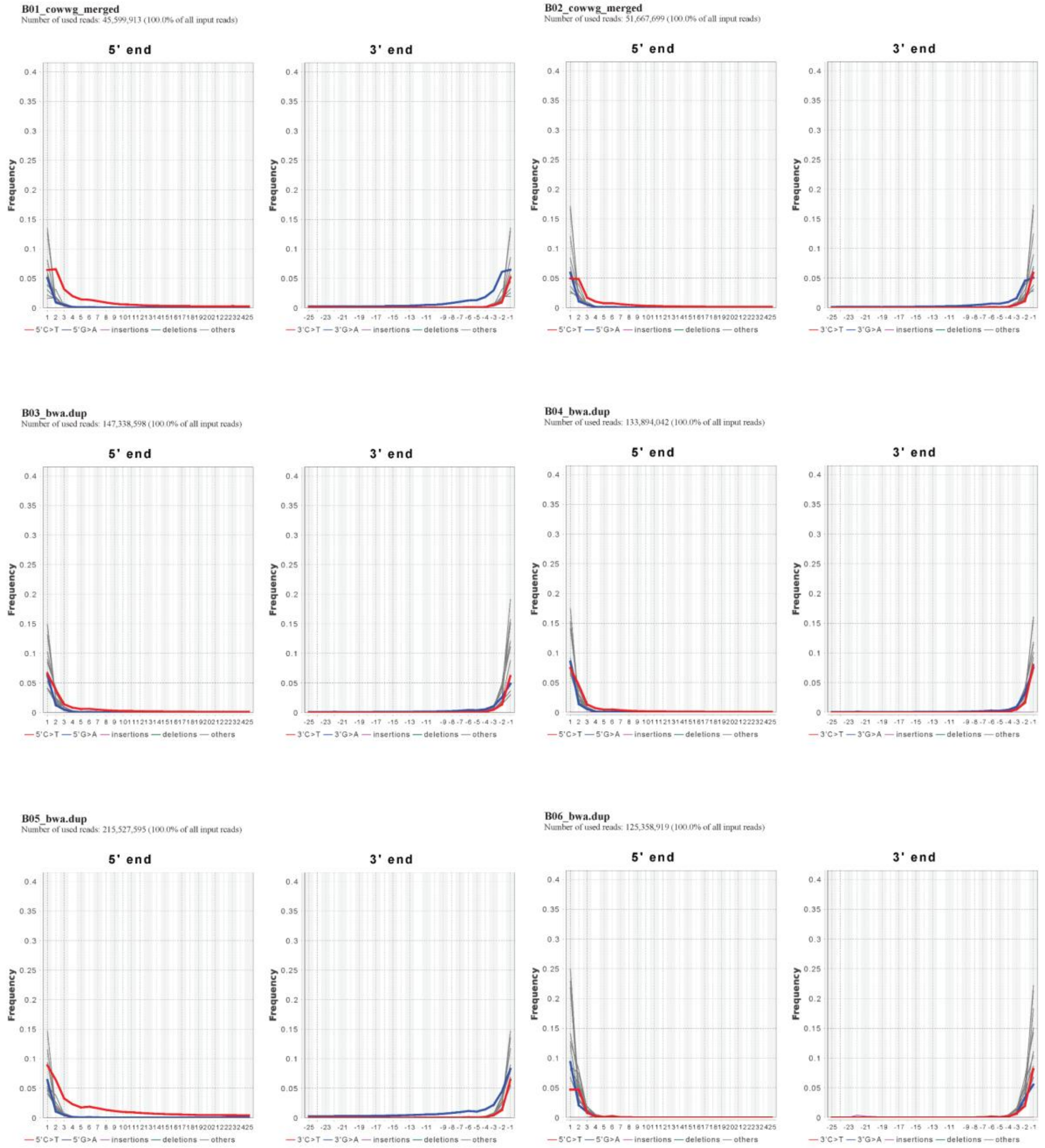


**Fig. S1.** Deamination patterns for ancient individuals produced using DamageProfiler.

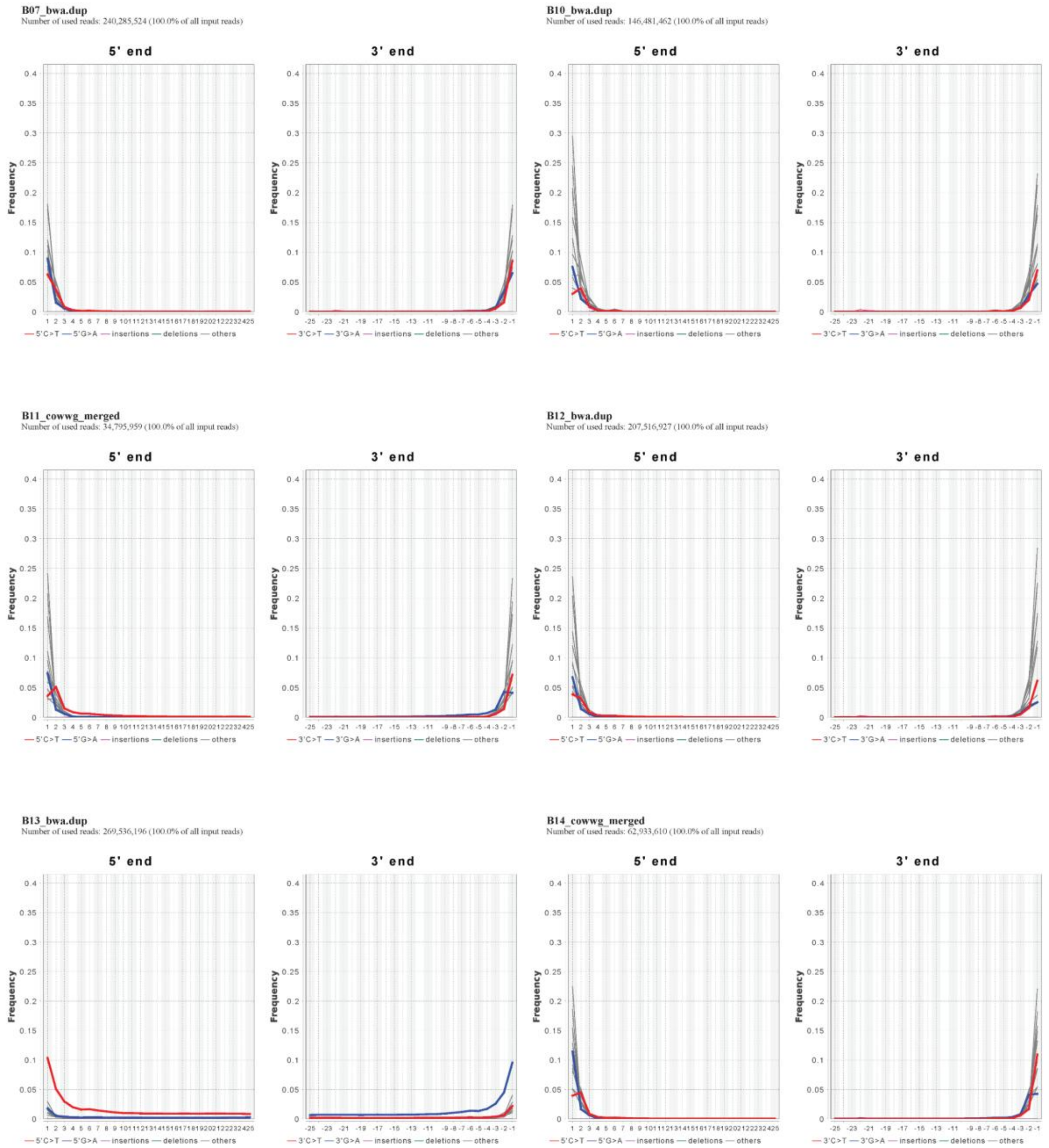


**Fig. S1 (continued).** Deamination patterns for ancient individuals produced using DamageProfiler.

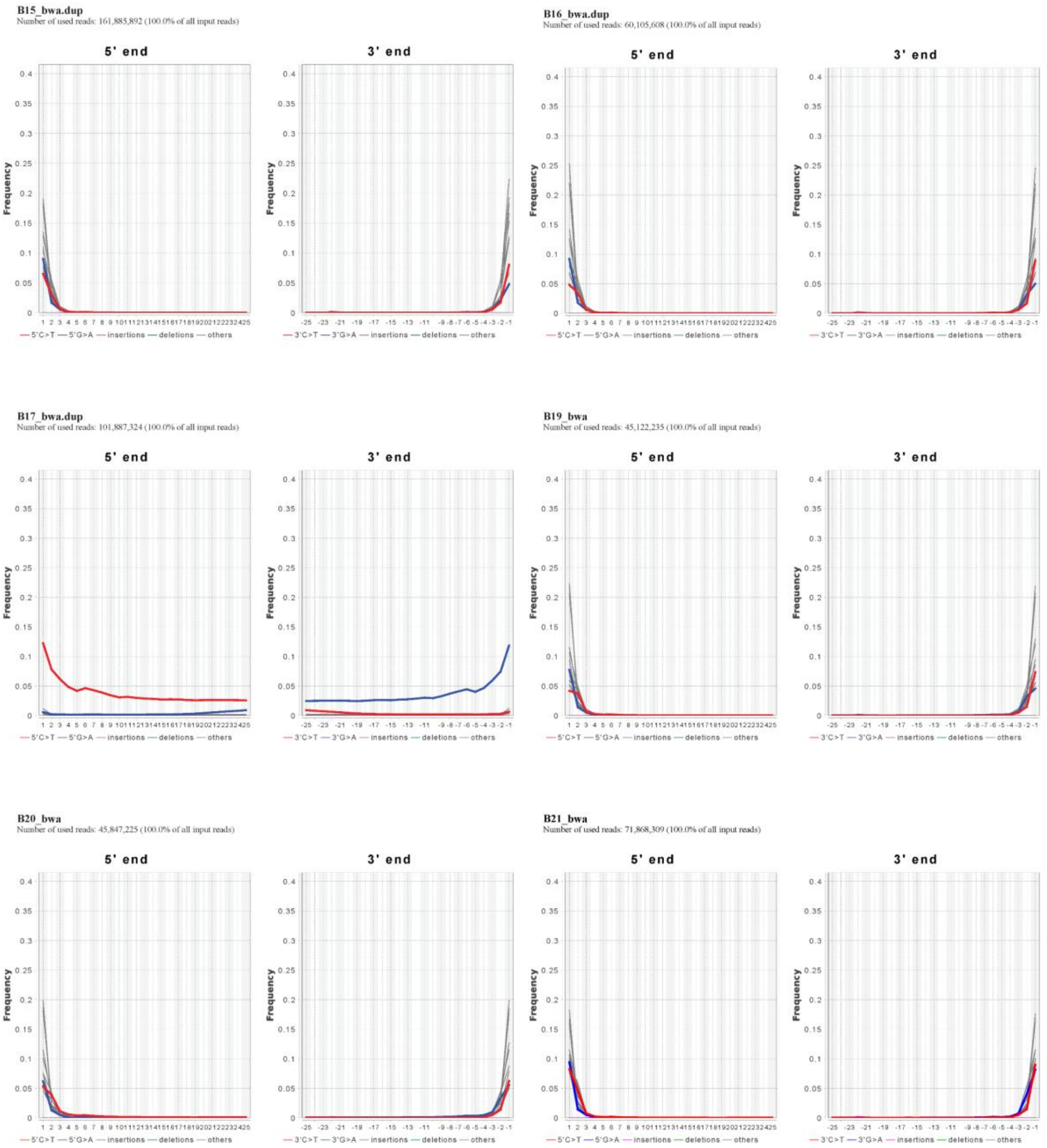


**Fig. S1 (continued).** Deamination patterns for ancient individuals produced using DamageProfiler.

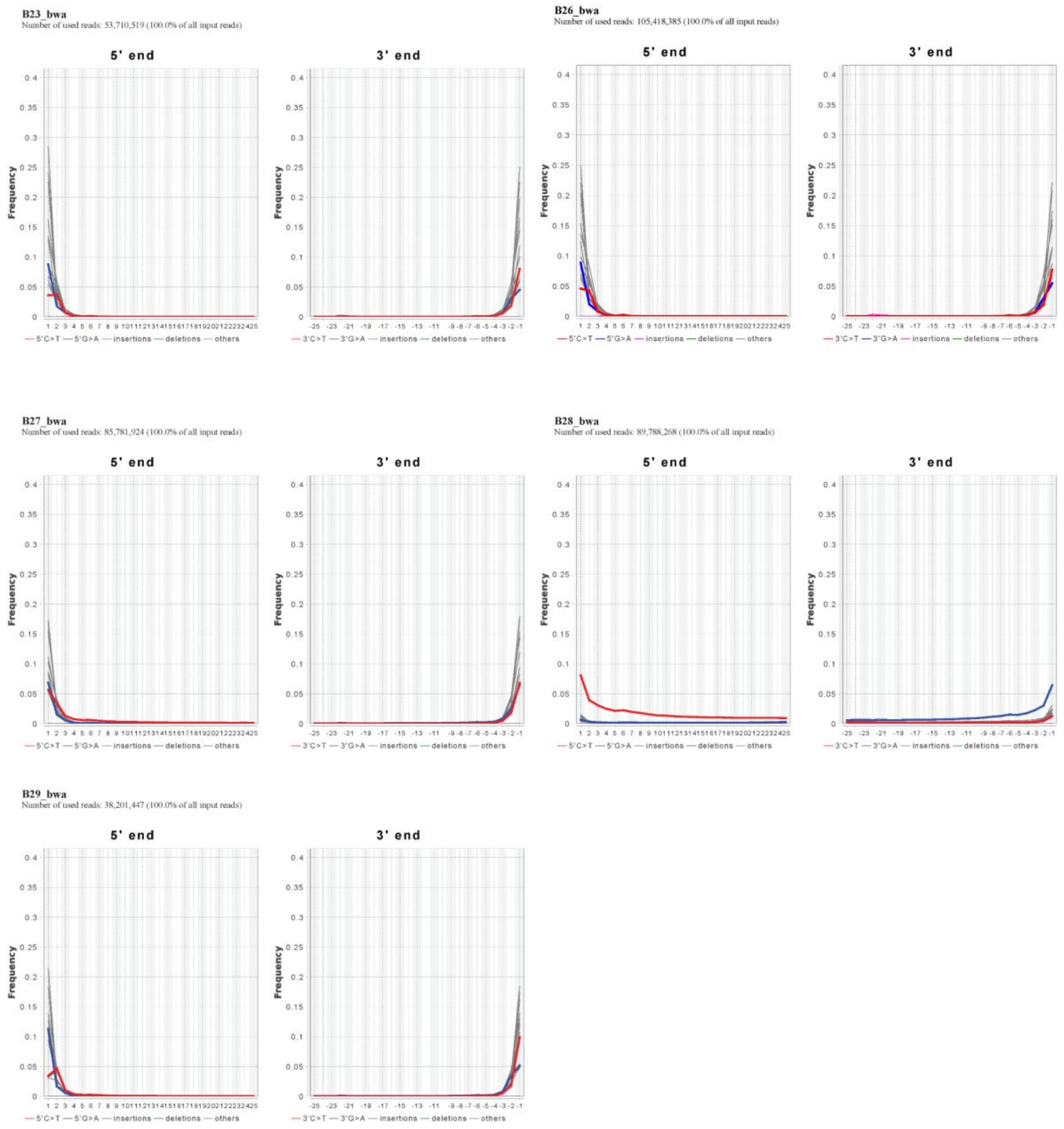


**Fig. S1 (continued).** Deamination patterns for ancient individuals produced using DamageProfiler.

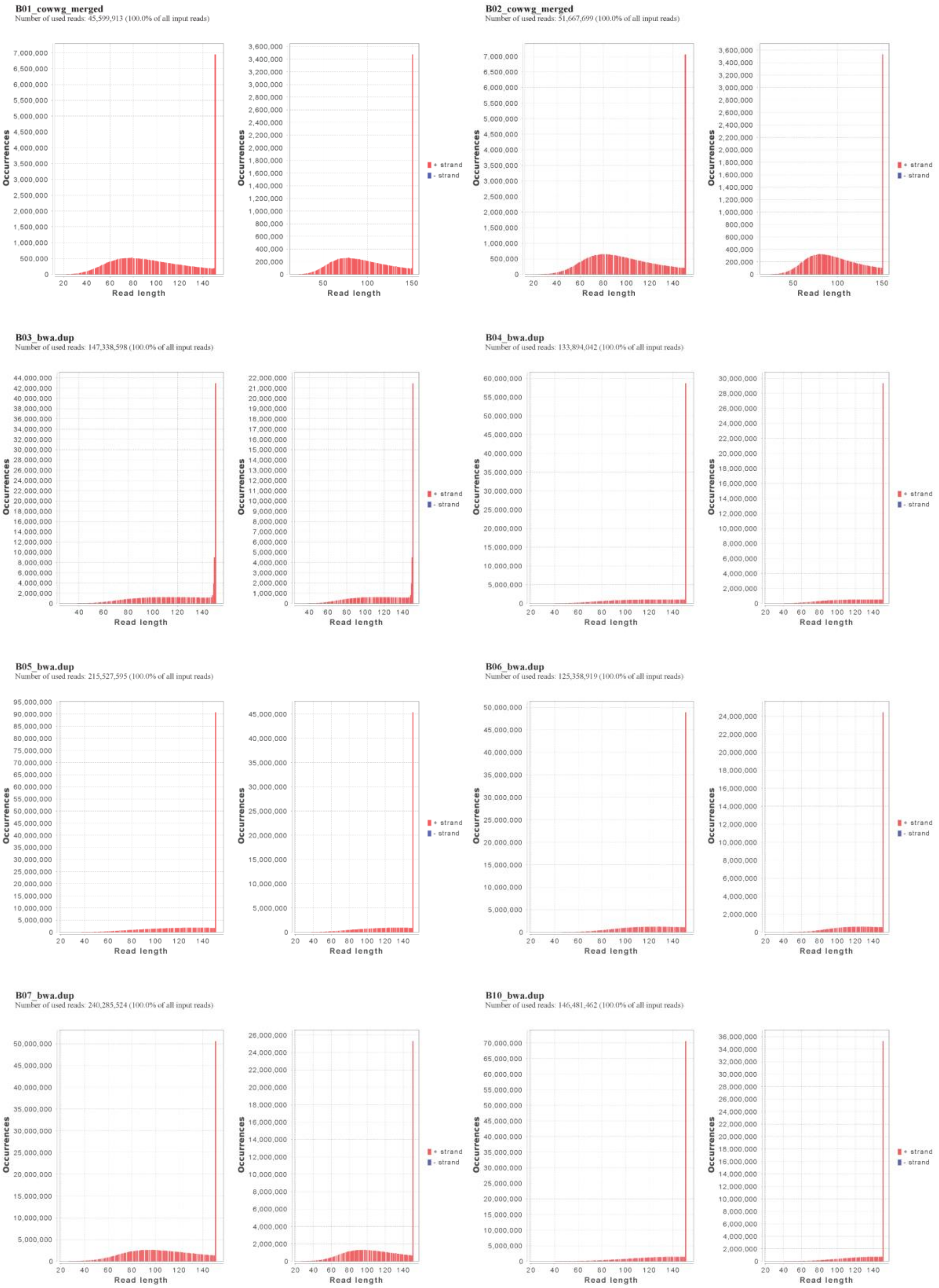


**Fig. S2.** Read length distributions for ancient individuals produced using DamageProfiler.

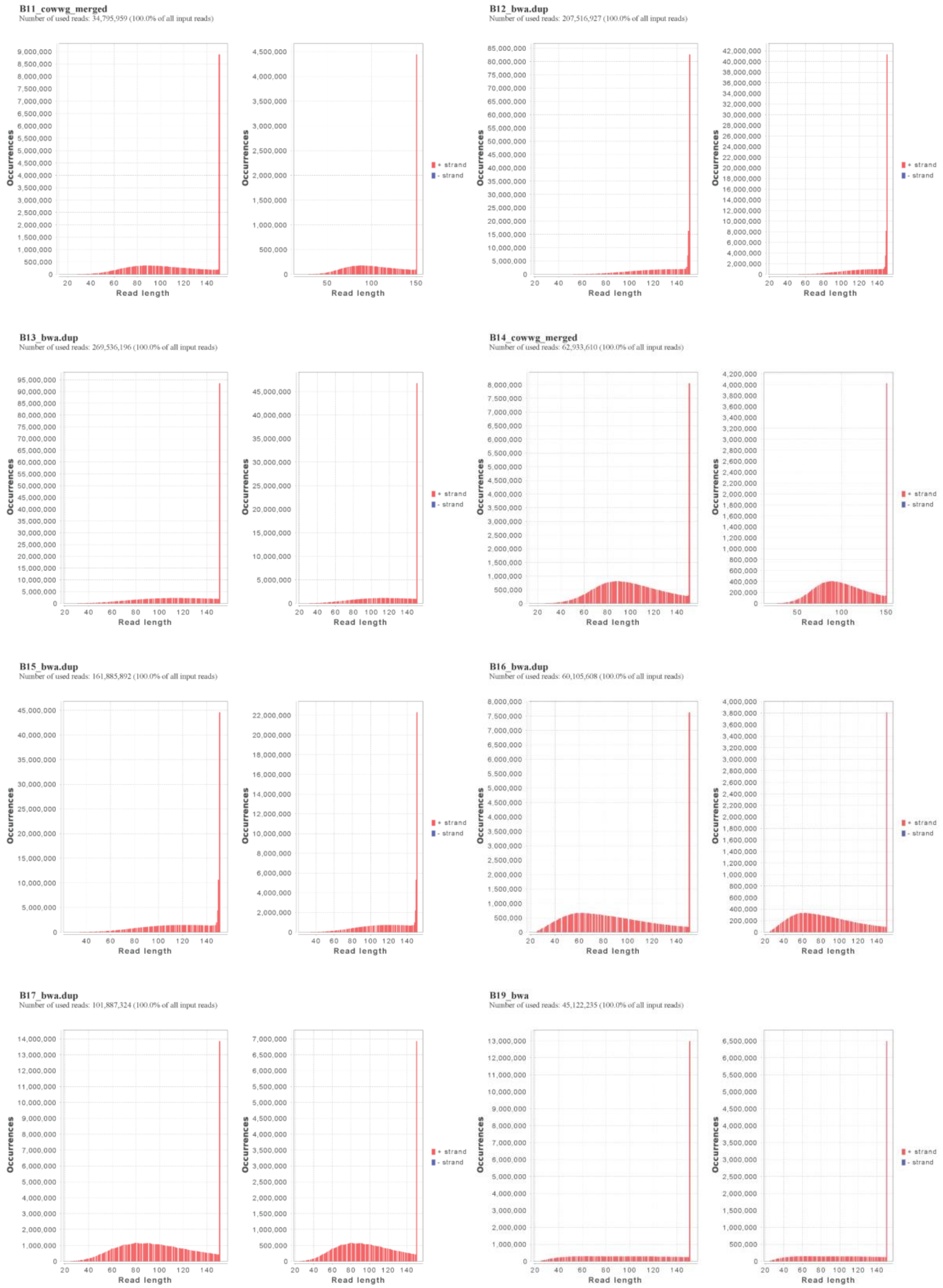


**Fig. S2 (continued).** Read length distributions for ancient individuals produced using DamageProfiler.

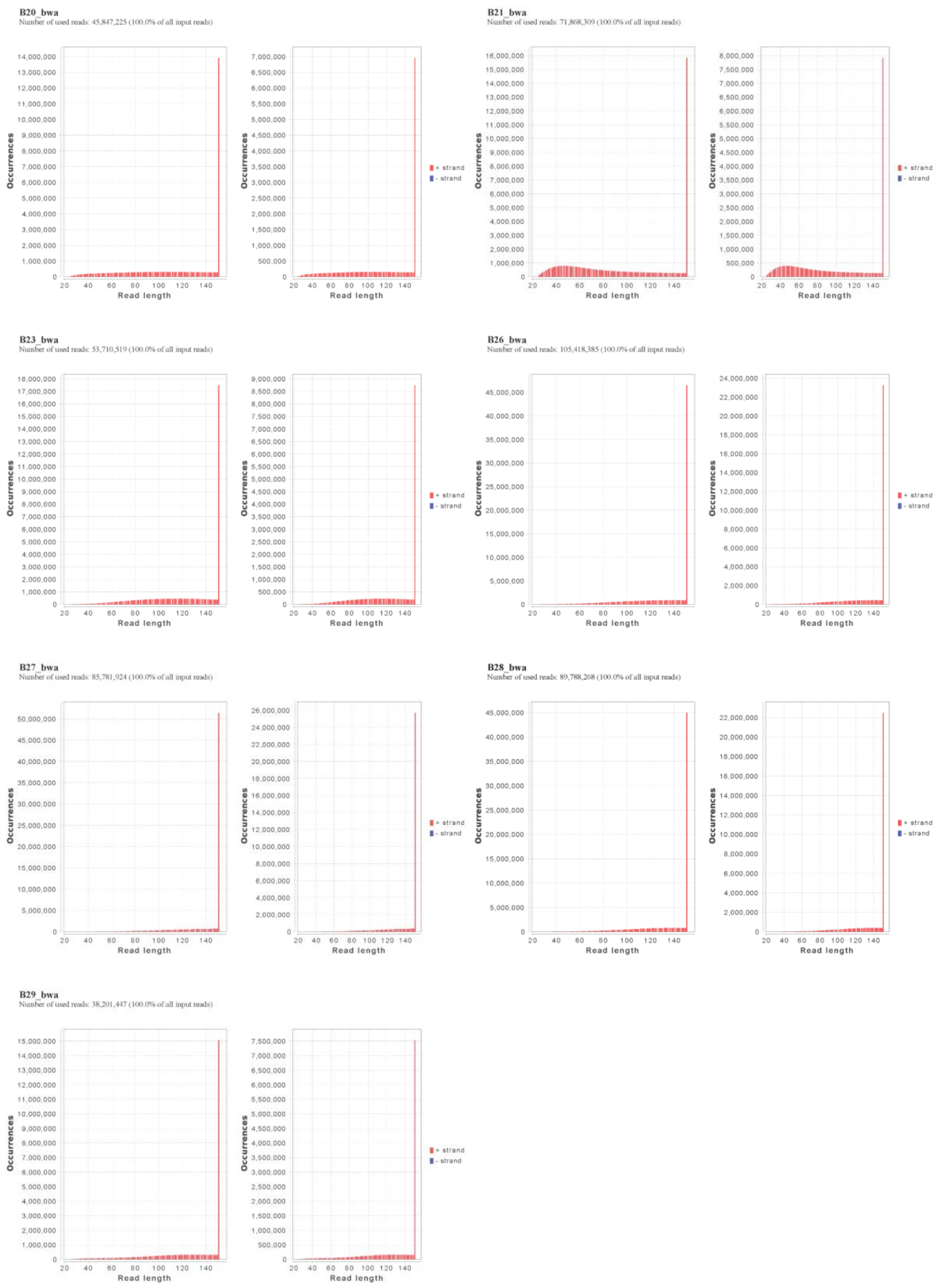


**Fig. S2 (continued).** Read length distributions for ancient individuals produced using DamageProfiler.

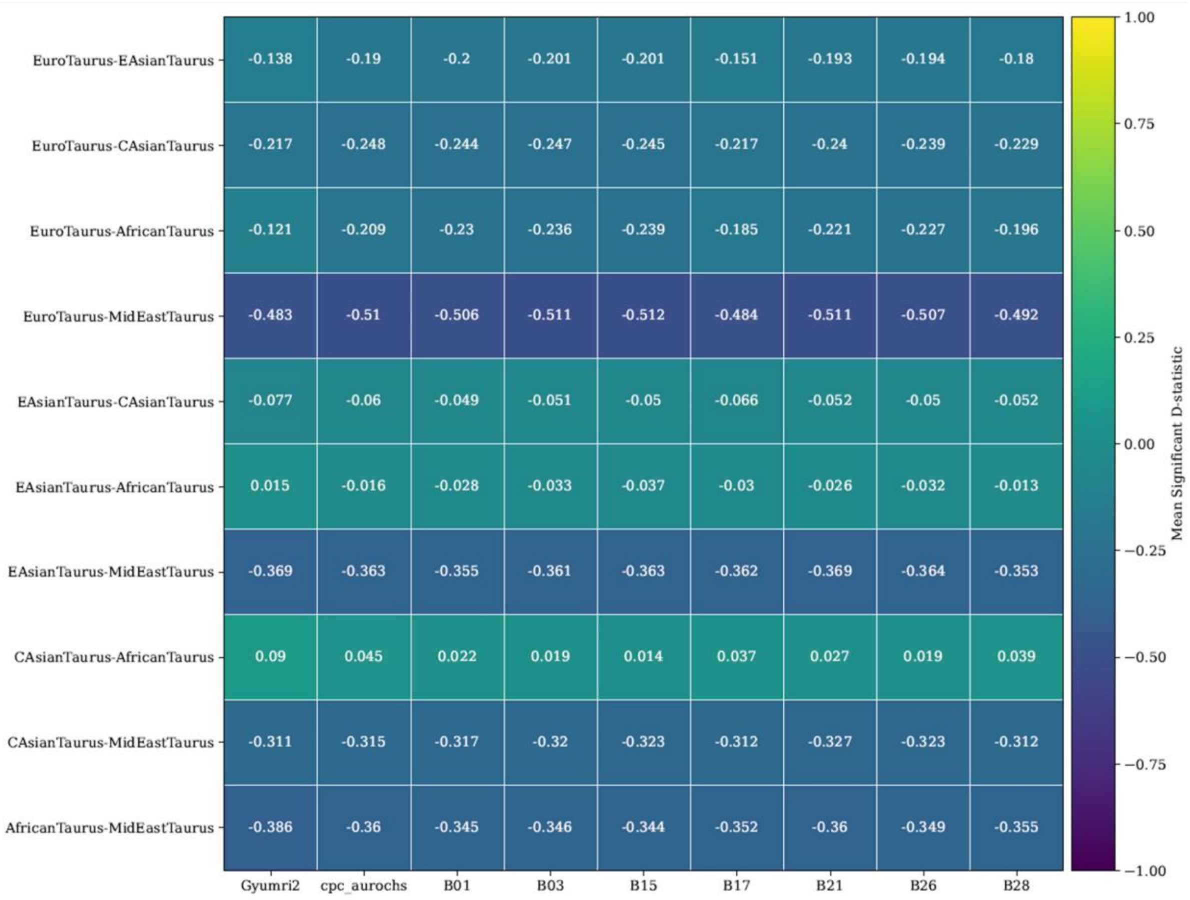


**Fig. S3.** D statistic heatmap for trios of taurine cattle.

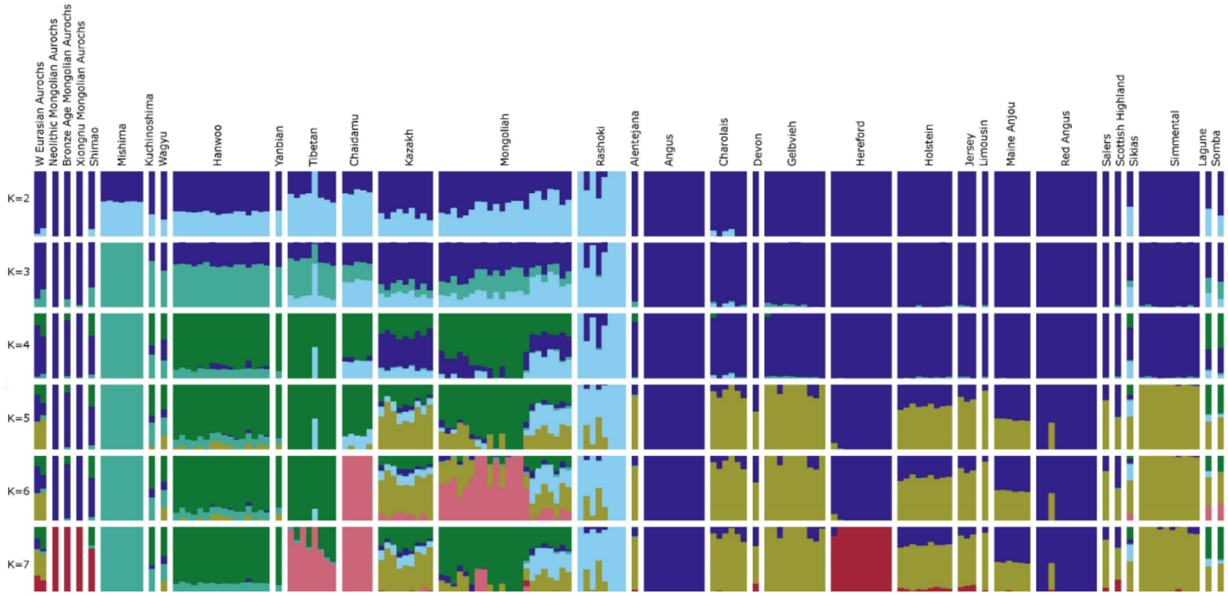


**Fig. S4.** Admixture graph showing aurochs and taurine cattle. Ancient Mongolian samples from three different time periods (Neolithic B01, Bronze Age B17, and Xiongnu B28) are included for comparison.

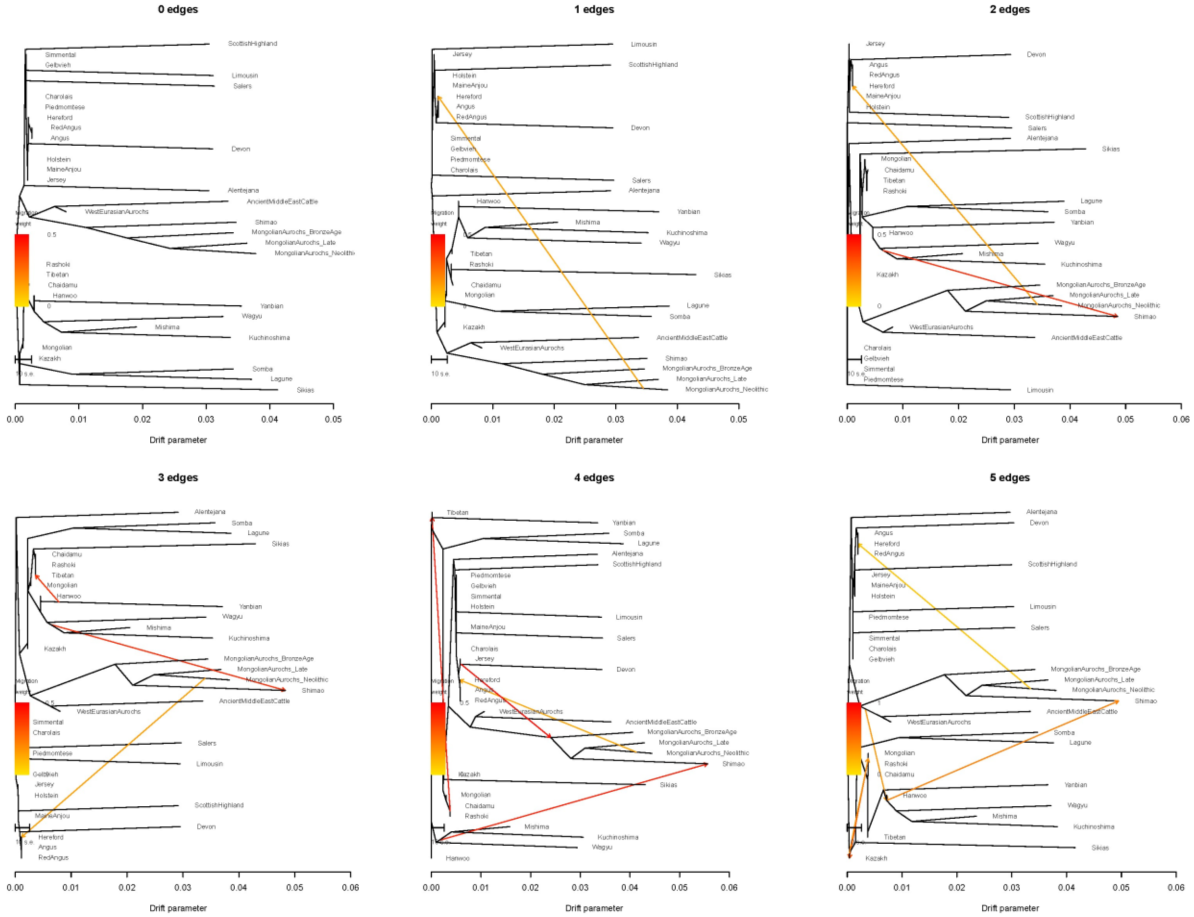


**Fig. S5.** Treemix plot showing 0-5 migration edges among taurine cattle populations. Here, B01 is labeled as MongolianAurochs_Neolithic, B17 is MongolianAuruchs_Bronze Age, and B28 is MongolianAurochs_Late.

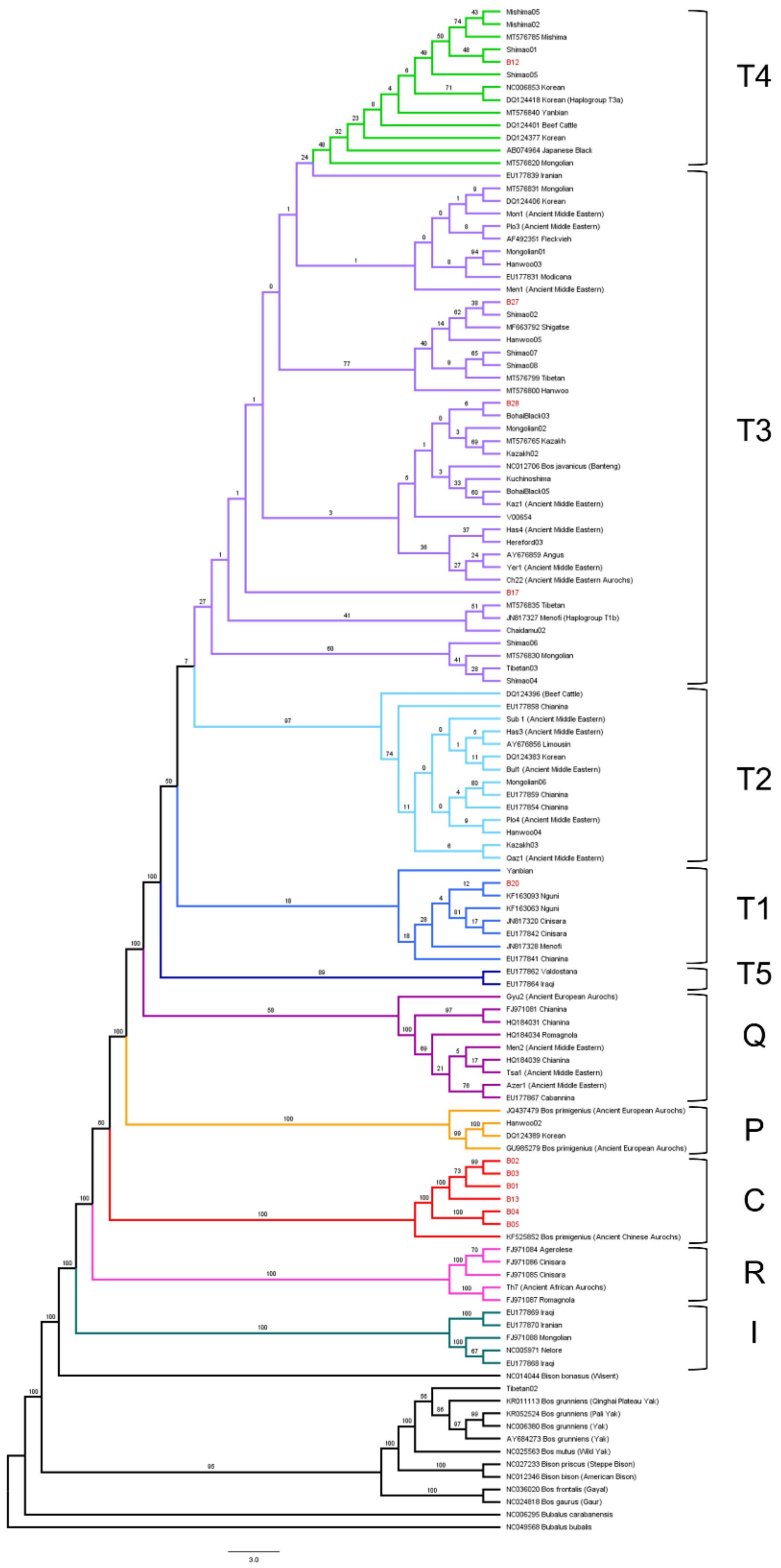

Mishima05
Mishima02
MT576785 Mishima
Shimao01
B12
Shimao05
NC006853 Korean
DQ124418 Korean (Haplogroup T3a)
MT576840 Yanbian
DQ124401 Beef Cattle
DQ124377 Korean
AB074964 Japanese Black
MT576820 Mongolian
T4
EU177839 Iranian
MT576831 Mongolian
DQ124406 Korean
Mon1 (Ancient Middle Eastern)
Plo3 (Ancient Middle Eastern)
AF492351 Fleckvieh
Mongolian01
Hanwoo03
EU177831 Modicana
Men1 (Ancient Middle Eastern)
B27
Shimao02
MF663792 Shigatse
Hanwoo05
Shimao07
Shimao08
MT576799 Tibetan
MT576800 Hanwoo
B28
BohaiBlack03
Mongolian02
MT576765 Kazakh
Kazakh02
NC012706 Bos javanicus (Banteng)
Kuchinoshima
BohaiBlack05
Kaz1 (Ancient Middle Eastern)
V00654
Has4 (Ancient Middle Eastern)
Hereford03
AY676859 Angus
Yer1 (Ancient Middle Eastern)
Ch22 (Ancient Middle Eastern Aurochs)
B17
MT576835 Tibetan
JN817327 Menofi (Haplogroup T1b)
Chaidamu02
Shimao06
MT576830 Mongolian
Tibetan03
Shimao04
T3
DQ124396 (Beef Cattle)
EU177858 Chianina
Sub 1 (Ancient Middle Eastern)
Has3 (Ancient Middle Eastern)
AY676856 Limousin
DQ124383 Korean
Bul1 (Ancient Middle Eastern)
Mongolian06
EU177859 Chianina
EU177854 Chianina
Plo4 (Ancient Middle Eastern)
Hanwoo04
Kazakh03
Qaz1 (Ancient Middle Eastern)
T2
Yanbian
B20
KF163093 Nguni
KF163063 Nguni
JN817320 Cinisara
EU177842 Cinisara
JN817328 Menofi
EU177841 Chianina
T1
EU177862 Valdostana
EU177864 Iraqi
T5
Gyu2 (Ancient European Aurochs)
FJ971081 Chianina
HQ184031 Chianina
HQ184034 Romagnola
Men2 (Ancient Middle Eastern)
HQ184039 Chianina
Tsa1 (Ancient Middle Eastern)
Azer1 (Ancient Middle Eastern)
EU177867 Cabannina
Q
JQ437479 Bos primigenius (Ancient European Aurochs)
Hanwoo02
DQ124389 Korean
GU985279 Bos primigenius (Ancient European Aurochs)
P
B02
B03
B01
B13
B04
B05
KF525852 Bos primigenius (Ancient Chinese Aurochs)
C
FJ971084 Agerolese
FJ971086 Cinisara
FJ971085 Cinisara
Th7 (Ancient African Aurochs)
FJ971087 Romagnola
R
EU177869 Iraqi
EU177870 Iranian
FJ971088 Mongolian
NC005971 Nelore
EU177868 Iraqi
I
NC014044 Bison bonasus (Wisent)
Tibetan02
KR011113 Bos grunniens (Qinghai Plateau Yak)
KR052524 Bos grunniens (Pali Yak)
NC006380 Bos grunniens (Yak)
AY684273 Bos grunniens (Yak)
NC025563 Bos mutus (Wild Yak)
NC027233 Bison priscus (Steppe Bison)
NC012346 Bison bison (American Bison)
NC036020 Bos frontalis (Gayal)
NC024818 Bos gaurus (Gaur)
NC006295 Bubalus carabanensis
NC049568 Bubalus bubalis
3.0

**Fig. S6.** Maximum likelihood tree of whole mitochondrial genome sequences of modern and ancient cattle and bovids. Samples are color-coded by mitochondrial haplogroup, which is labeled on the right. Ancient Mongolian sample names are highlighted in red.

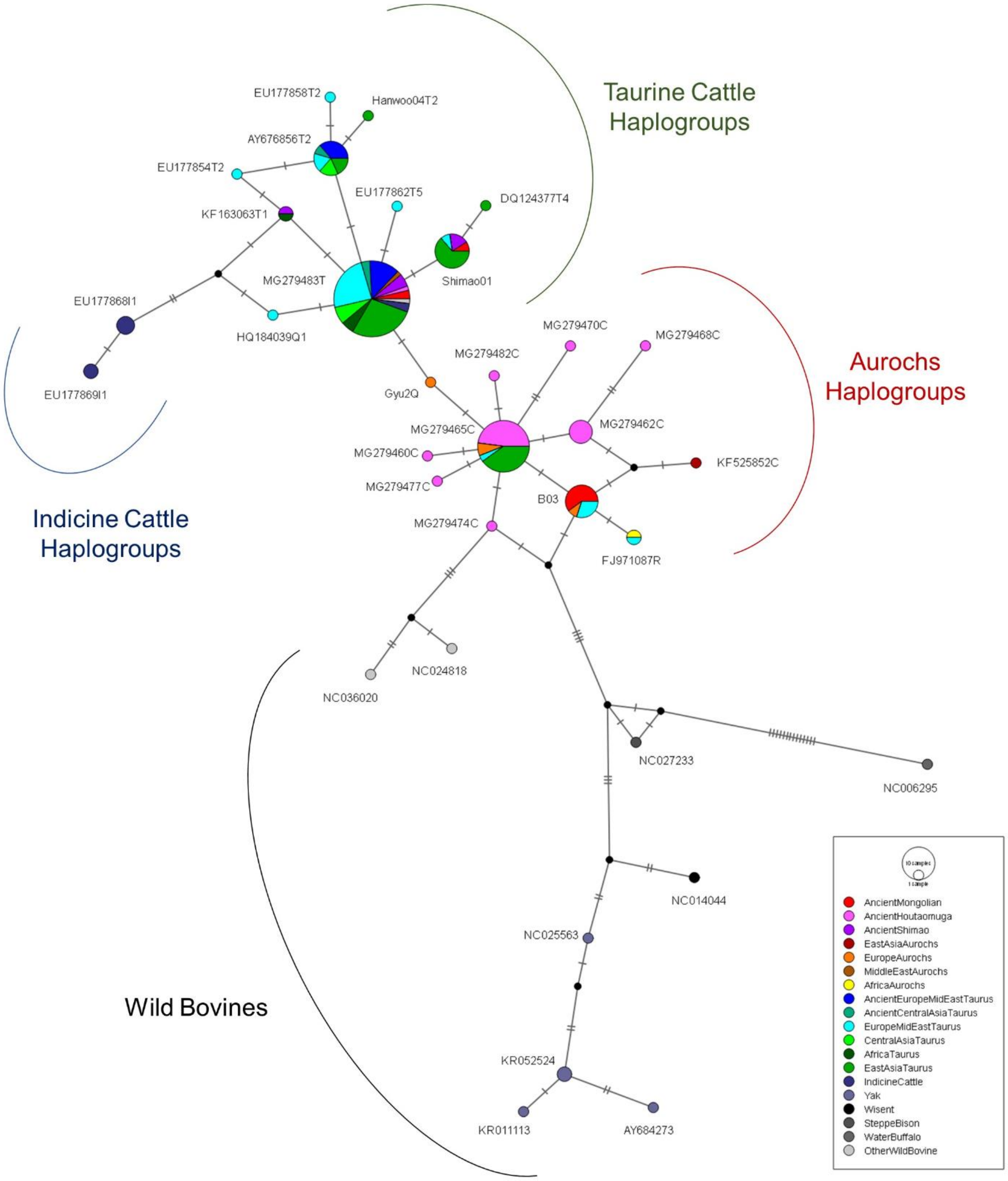

**Fig. S7.** Median joining network for the mitochondrial d-loop region. Ancient samples with low mtDNA coverage were not included in the analysis.

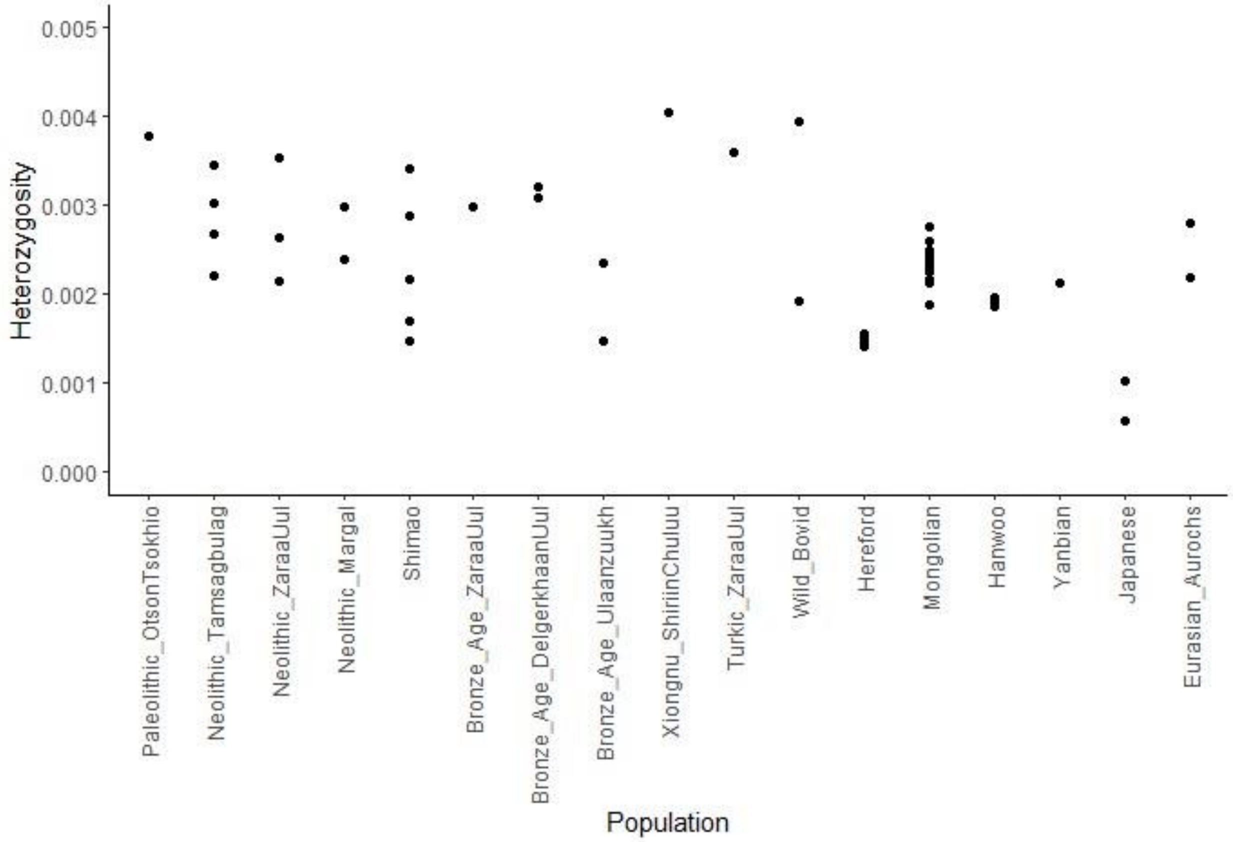


**Fig. S8.** Heterozygosity analysis of ancient Mongolian cattle genomes and comparative modern wild and domesticated bovines.

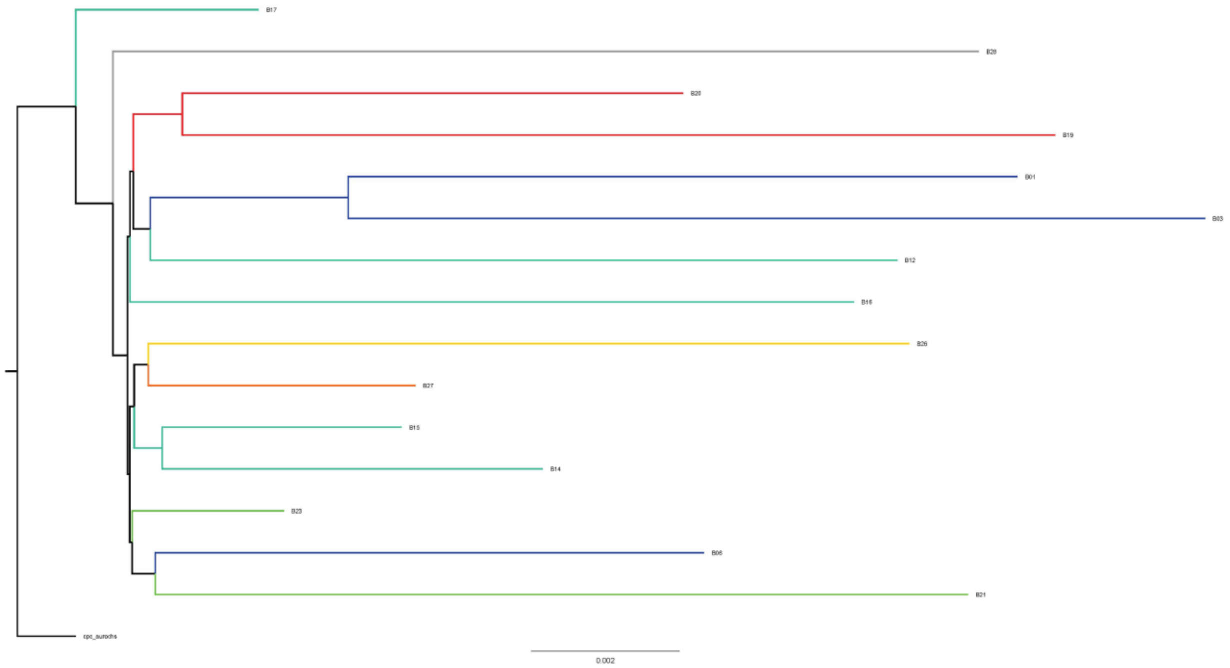


**Fig. S9.** Tree showing relationships between individual ancient Mongolian cattle genomes using only sites that are shared between all individuals. Individuals do not show any clear clustering by time period or region that would suggest changes in population structure through time, but B06 (Tamsagbulag), B15 (Zaraa Uul) and B23 (Margal) are all part of the same lineage and also show unusually high $\delta 13C$ and $\delta 15N$ values (**Table S1**). Yellow = Otson Tsokhio ; Dark blue = Tamsagbulag; Light blue = Zaraa Uul; Green = Margal; Red = Delgerkhaan Uul; Orange = Ulaanzuukh; Grey = Shiriin Chuluu.
